# Pan-cancer NET-associated immunothrombosis coupling identifies a neutrophil-linked but neutrophil-irreducible tumor ecology

**DOI:** 10.64898/2026.07.14.738403

**Authors:** Lin Zhang

## Abstract

**Background:** Neutrophil extracellular traps (NETs) connect innate immunity with thrombosis, metastasis and immune escape, but bulk-tumor NET-associated signatures may mainly report neutrophil abundance. We tested whether an established NETs score also captures a neutrophil-irreducible tumor ecology.

**Methods:** We reanalysed a published 19-gene NETs score in 9,496 TCGA tumors across 32 cancer types. Tumor-stratified correlations mapped Hallmark and custom immunothrombosis modules. Within each cancer type, the score was residualized against a curated neutrophil proxy and partitioned into four median-defined ecological quadrants. Clinical survival and two independent immune-checkpoint-blockade cohorts provided outcome context.

**Results:** Raw NETs score was positively associated with TNF-alpha/NF-kB signaling in 32/32 cancers (median Spearman rho=0.404; FDR<0.05 in 30) and hypoxia in 32/32 (median rho=0.394; FDR<0.05 in 31). After neutrophil residualization, hypoxia (median rho=0.206; positive in 30/32) and TNF-alpha/NF-kB signaling (median rho=0.165; positive in 29/32) remained dominant. NETs-high/neutrophil-low tumors retained myeloid-inflammatory, immunothrombotic and hypoxic enrichment. Residual NETs score was associated with adverse overall survival in eight of 23 evaluable cancers after available clinical adjustment. It did not discriminate response in IMvigor210 (AUC=0.519, P=0.642) or GSE91061 (AUC=0.527, P=0.808), although it differed across IMvigor210 immune phenotypes (P=0.045).

**Conclusions:** The published NETs score contains a neutrophil-linked yet incompletely neutrophil-reducible component that organizes hypoxic, inflammatory and immunothrombotic tumor ecology. Its translational value is context dependent rather than that of a universal standalone immunotherapy biomarker.

## Background

Neutrophil extracellular traps were first described as extracellular chromatin fibers decorated with antimicrobial proteins that immobilize and kill microorganisms [1]. The concept has since broadened from a discrete antimicrobial event to a family of chromatin-release programs with context-dependent roles in sterile inflammation, autoimmunity, vascular injury and cancer [2]. That breadth creates a measurement problem for human tumors: histological NET structures, circulating NET markers and bulk-tissue transcription are related but non-equivalent readouts. A transcriptomic NET-associated score should therefore be interpreted as an ecological proxy, not as direct proof that extracellular traps are physically present in every scored sample.

Cancer research nevertheless provides strong biological reasons to map this proxy. Contemporary reviews place NETs at the intersection of tumor initiation, dissemination, immune regulation and treatment failure [3]. Reciprocal tumor-neutrophil signaling can prime granulopoiesis and NET release, while NET-associated DNA, histones and proteases remodel local and distant tissue niches [4]. This bidirectionality suggests that a tumor-level score may contain information about the tissue state that induces NETs as well as the downstream effects of NET formation.

The immunothrombotic dimension is especially compelling. Tumor-bearing hosts can systemically prime neutrophils to release extracellular DNA that supports cancer-associated thrombosis [5]. NETs can also create adhesive vascular and parenchymal niches that facilitate metastatic seeding [6], and inflammatory NET remodeling can awaken dormant disseminated cancer cells [7]. These mechanisms imply that inflammation, coagulation, complement, endothelial activation and extracellular-matrix remodeling are not merely parallel correlates of NET-associated tumors; they may constitute a coupled host-defense program that malignant tissues exploit.

NETs also influence immune access. Tumor-derived CXCR1/CXCR2 agonists can induce traps that physically interfere with NK- and T-cell cytotoxicity [8]; IL-17-driven NETs can mediate checkpoint blockade resistance in pancreatic cancer [9]; and a tumor-derived Chi3l1-NET axis can promote T-cell exclusion in triple-negative breast cancer [10]. Recent work extends this logic to an NET-STC1 feedback circuit linked to immune evasion in bladder cancer [11] and to CCDC25-dependent epithelial-mesenchymal transition and treatment resistance in breast cancer [12]. Together, these studies predict that the biological meaning of a NET-associated transcriptional state depends on surrounding hypoxic, stromal and immune programs.

A 19-gene coefficient-weighted NETs score has already been reported as a pan-cancer prognostic signature [13]. The present study deliberately does not claim to develop another signature. Instead, it asks a different question that is essential for interpreting the published score: after the expected contribution of neutrophil abundance is modeled within each cancer type, does the remaining signal still organize a coherent tumor ecology? This distinction matters because many genes in NET-related signatures are expressed by neutrophils or myeloid cells. Without an abundance-aware analysis, a nominal NET signal can collapse into a relabeled leukocyte-content measure.

A second unresolved issue is the biological unit represented by a bulk-tumor score. NET formation is an event, whereas bulk RNA is a time-averaged mixture of malignant, immune, stromal and vascular cells. A score may therefore rise because more neutrophils are present, because resident neutrophils are activated, because other cells express overlapping inflammatory genes, or because the tissue environment favors NET formation. These explanations are not mutually exclusive, but they imply different experiments and clinical uses. An abundance-aware atlas cannot resolve every cellular source; it can, however, identify which ecological relationships survive a stringent first separation from estimated neutrophil quantity.

This reasoning generates three discriminating predictions. If the score is only an abundance marker, pathway coupling should largely disappear after neutrophil residualization and NETs-high/neutrophil-low tumors should resemble their NETs-low counterparts. If it reflects a broader tissue program, selected associations should persist in both analyses. Finally, if that program is a universal treatment biomarker, the same score should discriminate response across independent checkpoint-blockade cohorts. Testing these predictions in one workflow makes negative findings informative and prevents survival association alone from being used as proof of mechanism or clinical portability.

The Cancer Genome Atlas (TCGA) provides a harmonized foundation for this test [14]. Its pan-cancer immune landscape demonstrates that immune states recur across tissue lineages [15], whereas cell-of-origin structure remains a major determinant of molecular variation [16]. High-quality, uniformly curated survival endpoints permit tumor-stratified outcome analyses [17], and expression-derived stromal and immune estimates can provide admixture sensitivity checks where data are available [18]. These properties favor an analysis that standardizes and tests associations within each tumor type before summarizing cross-cancer consistency.

We therefore constructed an abundance-aware coupling atlas using selected Hallmark pathways from MSigDB [19] and transparent immune-cell modules informed by established tumor immune signatures [20]. We further examined the published score in pretreatment melanoma samples exposed to nivolumab [21] and in the IMvigor210 anti-PD-L1 urothelial cohort [22]. The prespecified organizing hypothesis was that NET-associated transcription would remain coupled to hypoxic and inflammatory programs after neutrophil adjustment, but would not function as a universally portable response classifier. The resulting design integrates raw coupling, within-cancer residualization, four-quadrant ecology, survival context and external immunotherapy evaluation.

## Methods

### Study design and data sources

This study was a retrospective, multi-cohort transcriptomic analysis. The discovery dataset comprised TCGA primary-tumor RNA expression, sample annotations and Pan-Cancer Clinical Data Resource outcomes. Samples were harmonized to 15-character TCGA sample barcodes and 12-character patient barcodes. Tumor types were retained when at least 30 samples had a finite NETs score. This yielded 9,496 tumor samples from 32 cancer types. Clinical analyses used one patient-level mean score when more than one tumor aliquot was available. No normal-tissue comparisons were used to support the principal conclusions.

External treatment context was evaluated in two independent cohorts: IMvigor210 urothelial cancer treated with atezolizumab and GSE91061 melanoma treated with nivolumab. Processed expression and clinical files already archived in the project were used. Pretreatment samples with mapped binary response labels were selected for response discrimination. All analyses used de-identified public data; no new participant recruitment, intervention or access to identifiable information occurred.

### Published NETs score reconstruction

The coefficient file accompanying the published 19-gene NETs prognostic signature was used without refitting. Gene symbols were harmonized, including IL8 to CXCL8, and the weighted score was computed as the sum of log-scale expression multiplied by the published coefficient for each matched gene. All 19 genes matched the TCGA and IMvigor210 expression matrices; 17 matched GSE91061 after Entrez-to-symbol conversion. Within each TCGA cancer type, the raw score was standardized to mean zero and unit variance; a global z score was used only for the cross-cancer distribution plot.

The source publication describes the variable as a NETs-related prognostic score. Existing project code and figures use the label NETosis. In this manuscript, NETs score and NET-associated score refer to the same expression-derived variable; neither label is intended to claim direct microscopy, citrullinated-histone measurement or biochemical quantification of extracellular traps.

### Pathway and immune-ecology modules

Thirteen prespecified Hallmark programs were selected to cover TNF-alpha/NF-kB signaling, hypoxia, inflammatory response, IL6/JAK/STAT3 signaling, complement, angiogenesis, coagulation, epithelial-mesenchymal transition, TGF-beta signaling, interferon-gamma response, IL2/STAT5 signaling, KRAS signaling and WNT/beta-catenin signaling. Five transparent custom modules represented a neutrophil core, immunothrombosis, myeloid inflammation, T-cell exhaustion/checkpoints and CAF-endothelial stroma. Module gene membership and matching diagnostics are supplied in the supplementary tables.

For each module, expression of every matched gene was z standardized across the analyzed expression matrix and the sample-level module score was the mean of those gene-wise z scores. Modules required at least three matched genes. The curated neutrophil-abundance proxy comprised MPO, ELANE, PADI4, S100A8, S100A9, CXCL8, MMP9, CTSG, LCN2, LTF, CEACAM8 and OLFM4; 10 of these 12 genes were available. This proxy was standardized within cancer type before residualization.

### Neutrophil residualization and quadrant ecology

Within each cancer type, ordinary least-squares regression modeled the standardized NETs score as a function of the standardized neutrophil proxy. The residual, termed neutrophil-residual NETs score, is the component linearly uncorrelated with the proxy in that cancer type. Residualization was used as a signal-separation device, not as evidence of biological independence or causality. Sensitivity residuals additionally included stromal score or tumor purity where available.

A complementary four-quadrant analysis dichotomized both the standardized NETs score and neutrophil proxy at their within-cancer medians. The key contrast compared NETs-high/neutrophil-low with NETs-low/neutrophil-low tumors, thereby holding the coarse neutrophil stratum constant. For each feature and cancer type, the effect was the difference in median module score between these groups. Wilcoxon rank-sum tests were applied when at least 10 finite observations and both groups were present.

### Correlation and cross-cancer consistency analyses

Spearman correlations were calculated separately within each cancer type between the raw or residual NETs score and each pathway module. A test required at least 10 complete observations and at least three unique values in both variables. Benjamini-Hochberg correction was performed across the 32 cancer-specific tests for each score-module combination. Cross-cancer consistency was summarized by the median rho, proportion of positive correlations and number of cancer types with false-discovery rate (FDR) below 0.05. The quadrant contrasts were analogously corrected within each feature across cancer types. All tests were two sided.

### Clinical outcome analyses

Overall survival (OS), disease-specific survival and progression-free interval were taken from the curated TCGA clinical outcome resource. Cox proportional-hazards models were fitted separately by cancer type when at least 50 patients and 20 events were available. The exposure was a one-standard-deviation increase in raw or neutrophil-residual NETs score. Univariable, age/sex-adjusted and age/sex/stage-adjusted model families were estimated. Covariates entered only when coverage and variation thresholds were met; thus the nominal fully adjusted family used age alone or age plus sex in cancers lacking usable stage. FDR correction was performed within endpoint, score and model family.

### Immunotherapy analyses

The published coefficients were applied to each external expression matrix after gene harmonization. Responders were complete or partial responders and nonresponders were stable or progressive disease, according to each processed clinical file. Wilcoxon tests compared standardized scores, and ROC AUC was oriented toward nonresponse. In IMvigor210, a Kruskal-Wallis test compared desert, excluded and inflamed immune phenotypes. These were external context analyses; no threshold was trained in TCGA and transferred to the treatment cohorts.

### Purity, genomic and reproducibility analyses

Expression-derived ESTIMATE immune, stromal and purity scores matched only 396 bladder cancer samples in the available local input. These analyses were therefore explicitly treated as a BLCA-only sensitivity assessment and not as pan-cancer purity correction. Available TMB, mutation-signature proxy and thresholded copy-number burden also mapped to only one cancer type and were reported solely as exploratory supplementary analyses. Neither restricted analysis was used to support a cross-cancer genomic or purity claim.

The complete workflow was rerun from the project entry point in R, followed by submission-asset finalization. Tables and figures were compared by SHA-256 hashes before and after rerun. OpenAI ChatGPT was used for code review, manuscript organization, English-language editing and reference cross-checking. All numerical analyses were executed locally in R; every reported number and reference was checked against the generated output or PubMed metadata and approved by the author. The language model was not treated as an author.

**Table 1.**
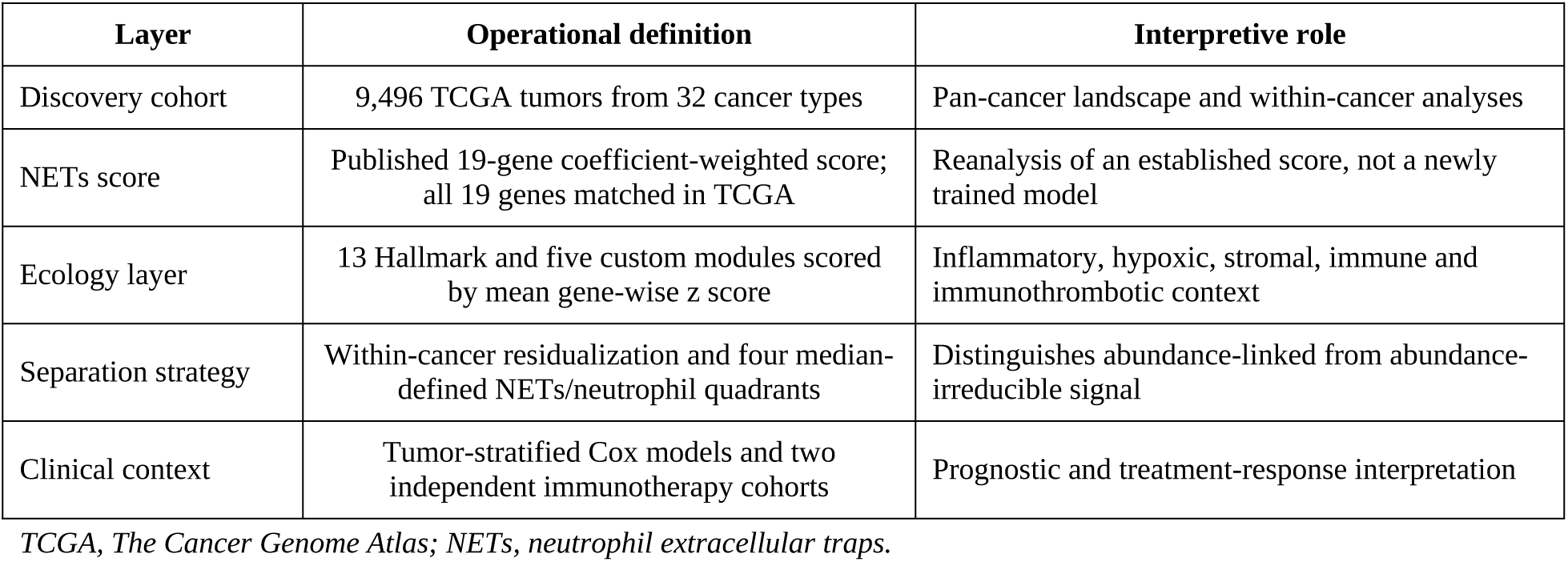
Study design and analytic layers.

## Results

### Cohort composition and pan-cancer distribution

The final discovery atlas included 9,496 tumors from 32 TCGA cancer types, ranging from 36 cholangiocarcinomas to 1,102 breast cancers. All 19 genes in the published coefficient set matched the TCGA expression matrix. Global standardization revealed marked lineage-level heterogeneity: glioblastoma, cholangiocarcinoma, esophageal carcinoma, lung squamous-cell carcinoma and hepatocellular carcinoma had the highest median global z scores. Because cell lineage and baseline immune composition differ substantially across tumors, all subsequent association tests used within-cancer standardized scores rather than pooling samples across cancer types (Fig. 1).

**Figure 1.**
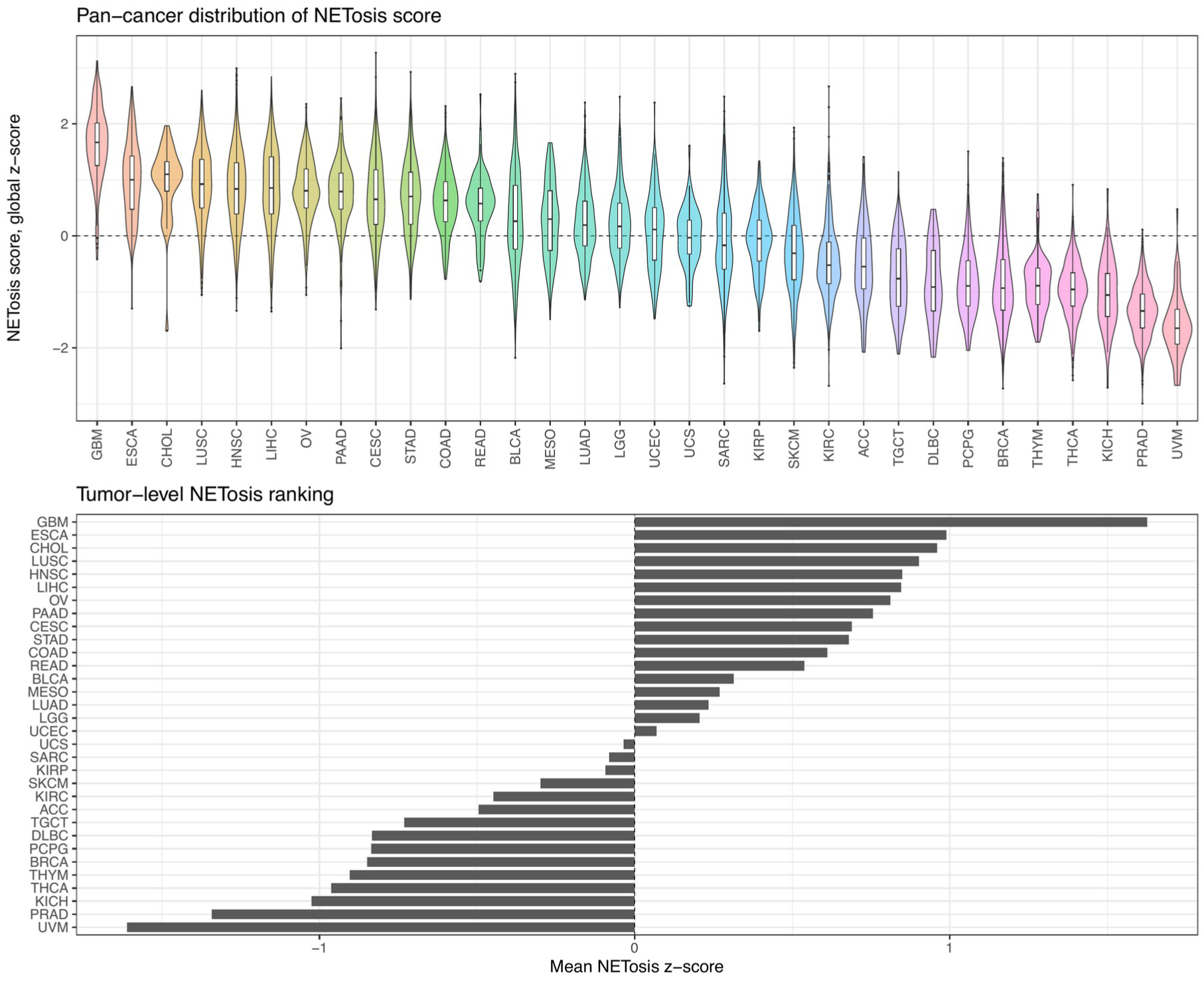
Pan-cancer distribution of the NETs score. Violin and box plots show the published coefficient-weighted NETs score across 9,496 TCGA tumors from 32 cancer types. Cancer types are ranked by median globally standardized score. Points and inner summaries display the observed sample distribution; downstream analyses used within-cancer standardization.

### Raw NETs score maps an inflammatory and immunothrombotic state

Tumor-stratified analyses showed a highly recurrent raw coupling pattern (Fig. 2; Table 2). TNF-alpha/NF-kB signaling was positively correlated with the NETs score in all 32 cancer types, with a median rho of 0.404 and FDR below 0.05 in 30. Hypoxia was likewise positive in all 32 cancers (median rho=0.394; FDR<0.05 in 31). The custom immunothrombosis core had a median rho of 0.278, was positive in 30 of 32 cancers and significant in 23. Inflammatory response, myeloid inflammation, IL6/JAK/STAT3, complement, angiogenesis and coagulation also had positive median correlations. These effects establish that the score is embedded in a broad tissue-stress and host-defense program, but they do not by themselves distinguish NET-associated activity from neutrophil content.

**Figure 2.**
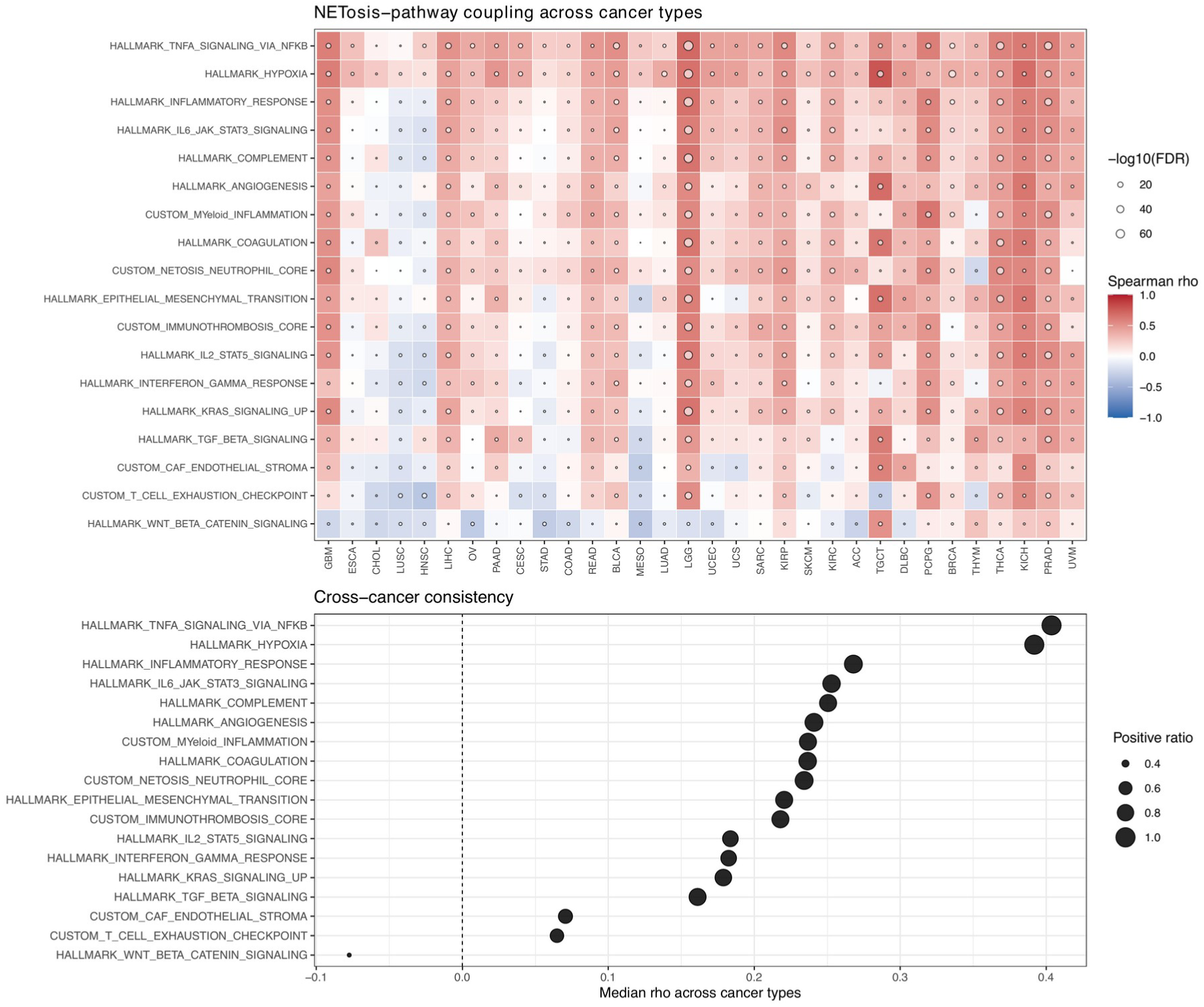
Raw NETs-pathway coupling across cancer types. The heat map reports within-cancer Spearman correlations between the standardized NETs score and selected Hallmark or custom modules. Color represents rho; point size represents -log10(FDR). The consistency panel summarizes median rho and the fraction of cancers with a positive association. FDR was controlled across cancer types for each score-module combination.

**Table 2.** Raw and residual pathway coupling.

| Score | Module | Median rho | Positive cancers | FDR<0.05 |
| --- | --- | --- | --- | --- |
| Raw NETs score | TNF-alpha Signaling Via NF-kB | 0.404 | 32/32 | 30 |
| Raw NETs score | Hypoxia | 0.394 | 32/32 | 31 |
| Raw NETs score | Immunothrombosis Core | 0.278 | 30/32 | 23 |
| Raw NETs score | Inflammatory Response | 0.272 | 29/32 | 25 |
| Neutrophil-residual score | Hypoxia | 0.206 | 30/32 | 20 |
| Neutrophil-residual score | TNF-alpha Signaling Via NF-kB | 0.165 | 29/32 | 17 |
| Neutrophil-residual score | TGF-beta Signaling | 0.078 | 22/32 | 17 |
| Neutrophil-residual score | Epithelial Mesenchymal Transition | 0.070 | 22/32 | 9 |
| Neutrophil-residual score | Immunothrombosis Core | 0.068 | 20/32 | 9 |

FDR, false-discovery rate. Rows are ordered by median rho within score type.

### Hypoxic and NF-kB coupling persists after neutrophil residualization

Residualization attenuated the magnitude of most associations, confirming that neutrophil abundance contributes substantially to the raw score. It did not erase the cross-cancer structure (Fig. 3). Hypoxia remained the leading residual correlate, with median rho=0.206, positive direction in 30 of 32 cancer types and FDR<0.05 in 20. TNF-alpha/NF-kB signaling ranked second (median rho=0.165; positive in 29 of 32; significant in 17). TGF-beta signaling, epithelial-mesenchymal transition and the immunothrombosis core retained smaller positive median associations (rho=0.078, 0.070 and 0.068, respectively). In contrast, checkpoint/exhaustion and CAF-endothelial stroma modules had negative median residual correlations. Thus the residual was not a weaker copy of the entire raw immune landscape; it selectively preserved hypoxic and inflammatory coupling while decoupling from several cell-content-associated modules.

**Figure 3.**
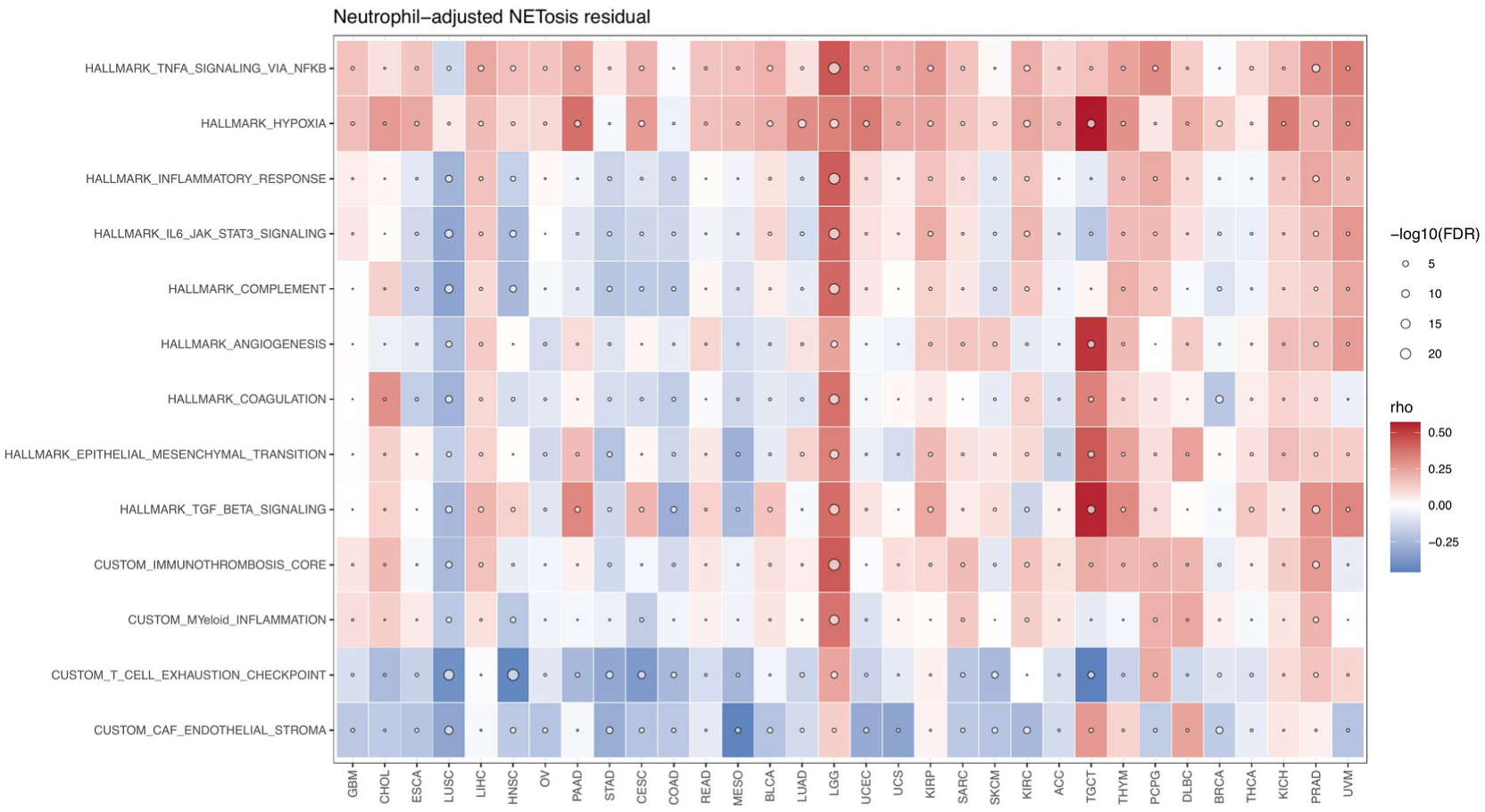
Neutrophil-adjusted residual NETs-pathway coupling. Within each cancer type, the standardized NETs score was regressed on a curated neutrophil-abundance proxy. The heat map displays Spearman correlations between the resulting residual and pathway modules. Color denotes rho and point size denotes -log10(FDR). Residualization is an operational signal-separation analysis and does not establish cell independence or causality.

### NETs-high neutrophil-low tumors retain immunothrombotic ecology

The four-quadrant analysis provided an assumption-light complement to linear residualization (Fig. 4). Because both axes were split at cancer-specific medians, every tumor type contained all four ecological states, although their fractions varied. Within the neutrophil-low stratum, NETs-high tumors showed the largest cross-cancer increase in myeloid inflammation (median delta=0.128; positive in 25 of 32 cancers; FDR<0.05 in eight), followed by immunothrombosis core (median delta=0.110; positive in 23; significant in eight) and TNF-alpha/NF-kB signaling (median delta=0.104; positive in 29; significant in 13). Hypoxia increased in 27 of 32 cancers and was significant in 16. The checkpoint/exhaustion and CAF-endothelial modules did not show the same positive pan-cancer shift. This contrast supports an ecologically specific NET-associated state among tumors with comparatively low neutrophil proxy values.

**Figure 4.**
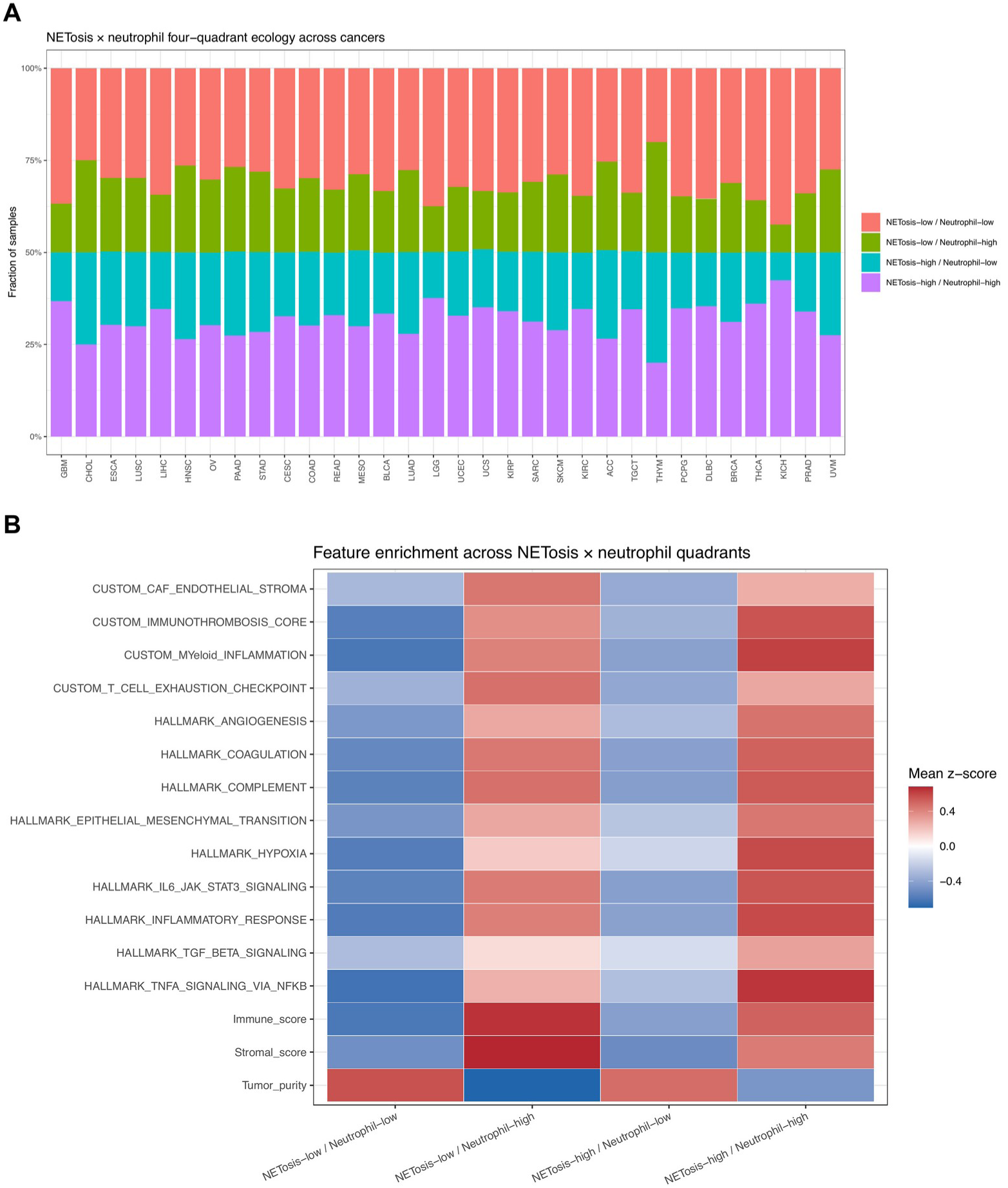
NETs-neutrophil ecological quadrants. (A) Fractions of samples in four cancer-specific median-defined quadrants. (B) Mean within-cancer z scores of pathway and ecology modules in each quadrant. The principal comparison contrasts NETs-high/neutrophil-low with NETs-low/neutrophil-low tumors, retaining the same coarse neutrophil stratum while varying the NETs score.

**Table 3.** NETs-high neutrophil-low quadrant contrasts.

| Module | Median delta | Positive cancers | FDR<0.05 |
| --- | --- | --- | --- |
| Myeloid Inflammation | 0.128 | 25/32 | 8 |
| Immunothrombosis Core | 0.110 | 23/32 | 8 |
| TNF-alpha Signaling Via NF-kB | 0.104 | 29/32 | 13 |
| Inflammatory Response | 0.073 | 23/32 | 9 |
| Angiogenesis | 0.065 | 23/32 | 7 |
| Hypoxia | 0.061 | 27/32 | 16 |
| Epithelial Mesenchymal Transition | 0.060 | 25/32 | 9 |
| IL6 JAK STAT3 Signaling | 0.050 | 22/32 | 5 |

*Delta is the median-score difference between NETs-high/neutrophil-low and NETs-low/neutrophil-low tumors.*

### Residual NETs score has context-dependent survival associations

Twenty-three cancer types met the OS modeling criteria. In the nominal age/sex/stage model family, higher neutrophil-residual NETs score was associated with shorter OS at FDR<0.05 in eight cancers: LGG, KIRC, UVM, PAAD, LUAD, KIRP, BRCA and CESC (Fig. 5; Table 4). The strongest association was in LGG (HR=1.80 per standard deviation, 95% CI 1.44-2.26, FDR=7.85e-6), followed by UVM (HR=2.67, 95% CI 1.42-5.01, FDR=0.014) and KIRP (HR=1.91, 95% CI 1.23-2.97, FDR=0.015). KIRC, PAAD and LUAD showed more moderate but significant effects. Covariate availability varied by tumor type, and the adjusted model family used only available eligible covariates. The heterogeneity of effect sizes and evaluable covariates argues against presenting the residual score as a universal prognostic instrument.

**Figure 5.**
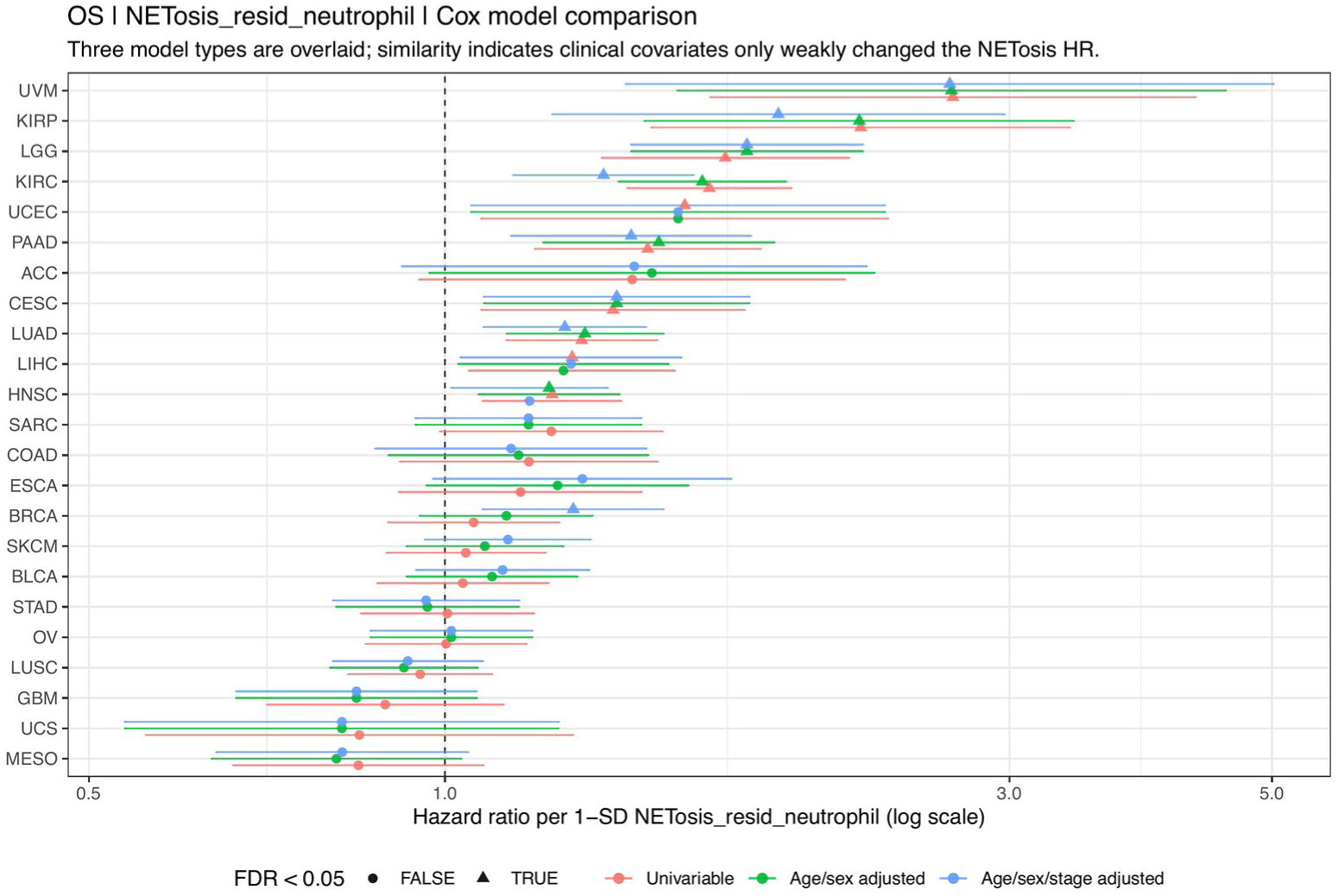
Overall-survival associations of residual NETs. Forest plots show hazard ratios per one-standard-deviation increase in neutrophil-residual NETs score for univariable, age/sex-adjusted and age/sex/stage-adjusted model families. Points indicate estimates, horizontal lines indicate 95% confidence intervals and filled shapes indicate FDR<0.05. Actual covariates used depended on tumor-specific coverage and variation.

**Table 4.**
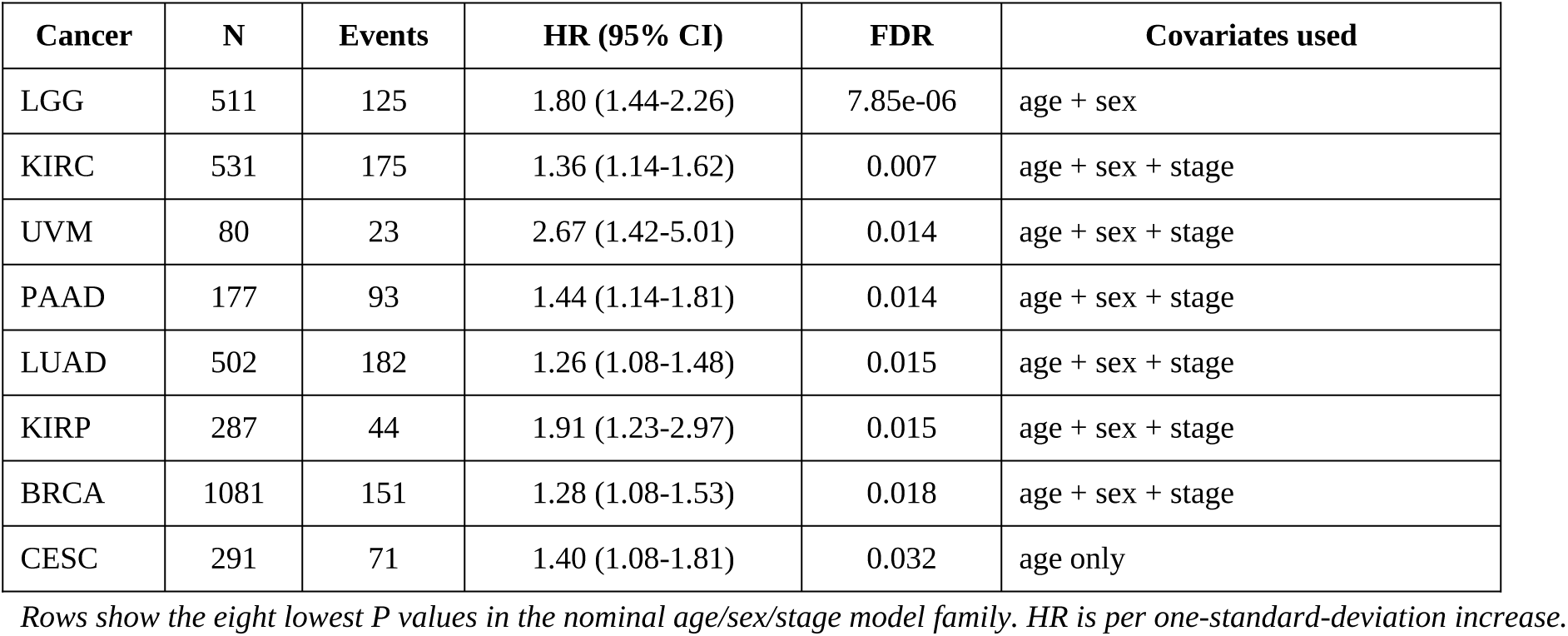
Residual NETs overall-survival associations.

### Checkpoint coupling does not translate into universal response discrimination

Raw NETs score showed heterogeneous correlations with immune-checkpoint genes across TCGA cancers (Fig. 6A), prompting direct external evaluation rather than inference from checkpoint expression alone. In IMvigor210, all 19 signature genes matched and 298 patients had binary response data (68 responders, 230 nonresponders). Median score was 0.174 standard-deviation units higher in nonresponders, but the difference was not significant (P=0.642) and discrimination was near chance (AUC=0.519; Fig. 6B). In GSE91061, 17 genes matched and 47 pretreatment samples had mapped outcomes (10 responders, 37 nonresponders). The corresponding difference was 0.231, P=0.808 and AUC=0.527 (Fig. 6C). In contrast, score distributions differed across IMvigor210 immune phenotypes (n=284; Kruskal-Wallis P=0.045), with medians of −0.219, −0.044 and 0.077 in desert, excluded and inflamed tumors. The score therefore tracked immune context weakly but was not a standalone response classifier.

**Figure 6.**
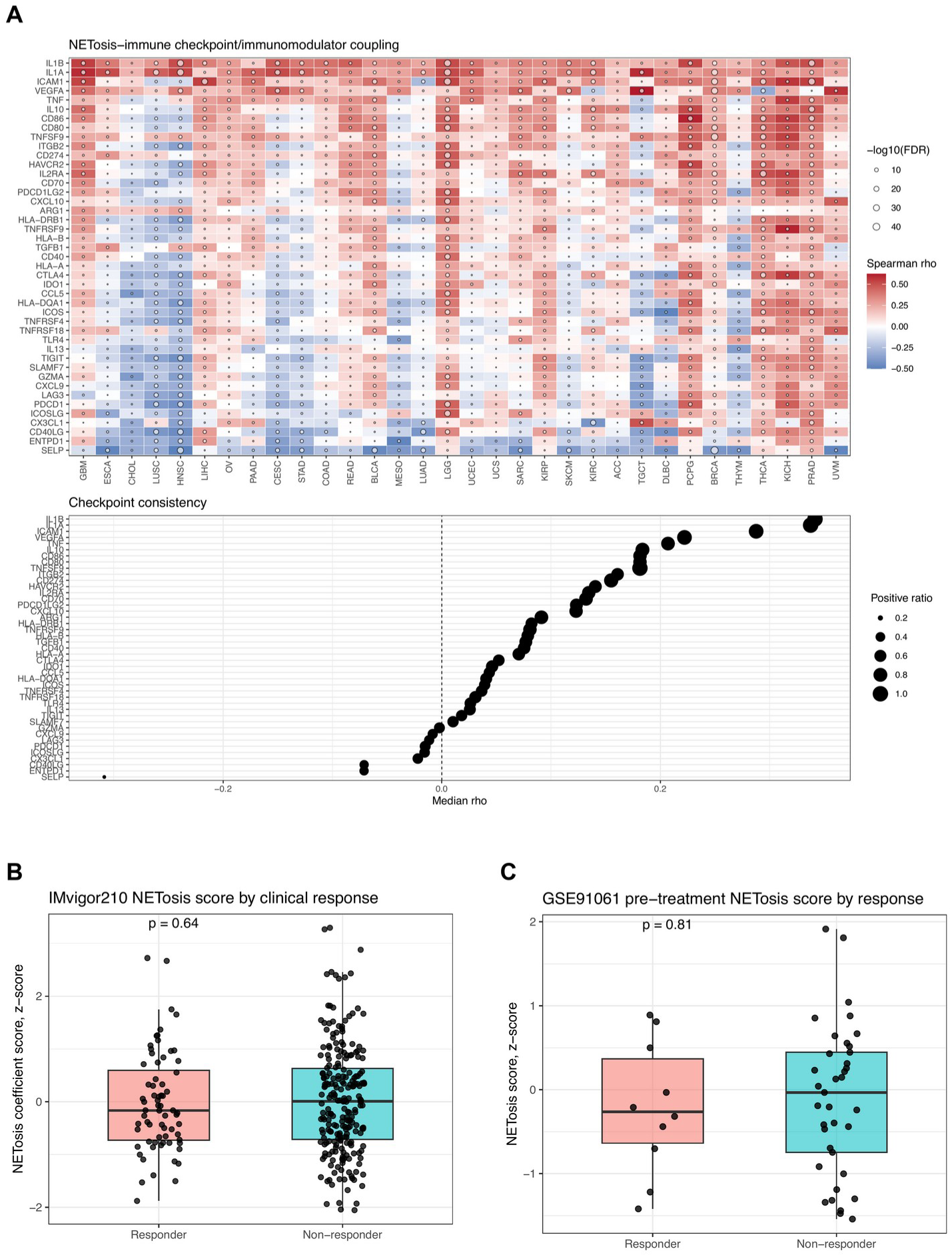
Checkpoint coupling and immunotherapy context. (A) Cancer-specific Spearman correlations between the NETs score and selected immune-checkpoint genes in TCGA. (B) Standardized score by binary response in IMvigor210 urothelial cancer. (C) Pretreatment score by binary response in GSE91061 melanoma. Box plots show medians and interquartile ranges; P values are two-sided Wilcoxon tests.

**Table 5.**
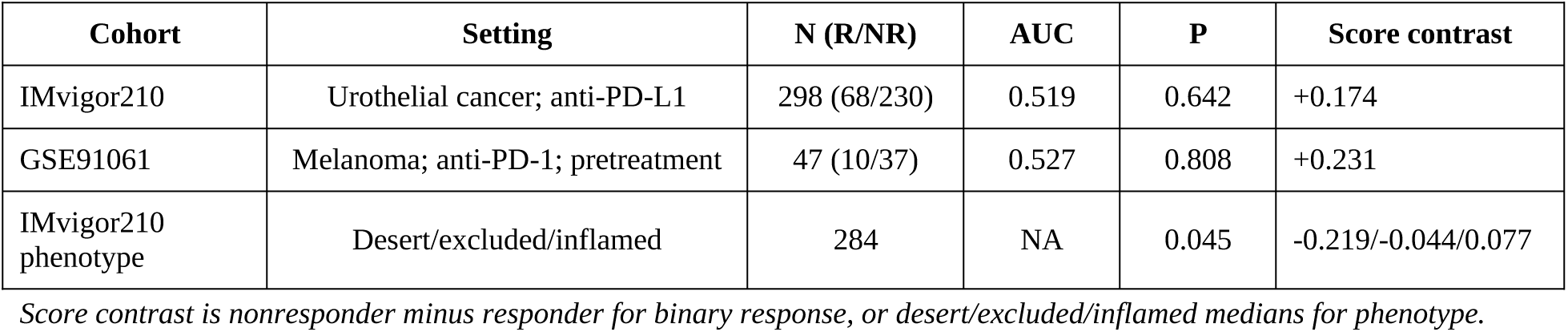
Immunotherapy cohort context.

| Cohort | Setting | N (R/NR) | AUC | P | Score contrast |
| --- | --- | --- | --- | --- | --- |
| IMvigor210 | Urothelial cancer; anti-PD-L1 | 298 (68/230) | 0.519 | 0.642 | +0.174 |
| GSE91061 | Melanoma; anti-PD-1; pretreatment | 47 (10/37) | 0.527 | 0.808 | +0.231 |
| IMvigor210 phenotype | Desert/excluded/inflamed | 284 | NA | 0.045 | -0.219/-0.044/0.077 |

*Score contrast is nonresponder minus responder for binary response, or desert/excluded/inflamed medians for phenotype.*

### Restricted sensitivity analyses define the boundary of inference

The available ESTIMATE file matched 396 BLCA samples only. In that restricted subset, raw NETs score correlated positively with immune score (rho=0.249), stromal score (rho=0.134) and neutrophil proxy (rho=0.460), and negatively with tumor purity (rho=-0.200; all FDR<0.01). After additional adjustment for stromal score or purity within BLCA, hypoxia, TNF-alpha/NF-kB and immunothrombosis correlations remained positive (Supplementary Fig. S1). However, these data cannot establish pan-cancer purity robustness. Genomic features likewise covered one tumor type; correlations with TMB, mutation-signature proxy and absolute thresholded CNV burden were small and are shown only in Supplementary Fig. S2. These restricted analyses were retained to make data limitations visible, not to enlarge the principal claim.

## Discussion

This study changes the interpretation of an established NETs prognostic score rather than proposing a new one. The earlier pan-cancer study demonstrated that a 19-gene model could stratify outcome. Our analysis shows what that score appears to represent at the tumor-ecology level: a large neutrophil-linked component plus a smaller, reproducible residual aligned most strongly with hypoxia and TNF-alpha/NF-kB signaling. The four-quadrant analysis reaches the same conclusion without depending on linear residuals. Together, these results define the central advance: the NET-associated signal is abundance linked, but not exhausted by a curated neutrophil-abundance proxy. We use ‘neutrophil-irreducible’ in this operational sense, not to imply that the residual is generated outside neutrophils.

Three linked design choices make that interpretation more than a conventional signature correlation study. First, every association was estimated within cancer type, preventing lineage-level differences from driving the result. Second, residualization directly tested the dependence of the score on a prespecified neutrophil proxy instead of treating neutrophil enrichment as an after-the-fact annotation. Third, the median-defined quadrants reproduced the key pattern without assuming a linear relation between the two scores. Convergence between these analytically distinct views is the strongest evidence in the study: raw coupling identifies the full NET-associated ecosystem, whereas residual and quadrant analyses isolate the part least compatible with a simple abundance explanation.

Cross-cancer consistency is important here, but uniformity is neither expected nor desirable. Cancer types differ in lineage, vascular architecture, treatment history, mutational landscape and baseline leukocyte composition. We therefore summarized direction and median effect rather than fitting one pooled pan-cancer coefficient. A pathway that is positive in most cancers yet strong in only a subset may represent a shared ecological tendency whose clinical consequences depend on local constraints. Conversely, tumor-specific outliers can reveal settings in which the 19-gene score is driven by alternative cell sources or biology. The atlas is consequently a map of reproducibility and heterogeneity, not a claim that all tumors instantiate one NET program.

This distinction is biologically plausible because tumor-associated neutrophils are heterogeneous in developmental state, localization and function [23]. Their phenotypes can range from direct tumor control to trophic, angiogenic and immunosuppressive programs, and cell number alone cannot encode that functional diversity [24]. A weighted score containing inflammatory receptors, chemotactic genes, tissue factor and degranulation-associated genes may therefore respond to activation state, neighboring-cell expression and tissue stress in addition to neutrophil quantity. Residualization does not identify the cellular source, but it helps prevent abundance from being the only explanation.

The residual should consequently be read as a contrast, not as a purified molecular entity. A positive residual can arise when the weighted 19-gene program is higher than expected for the measured neutrophil proxy. That mismatch may reflect activated neutrophils, spatially concentrated NET-producing cells, expression of score genes by tumor or stromal compartments, or imperfect sensitivity of the proxy. The consistent association with hypoxia and NF-kB narrows these possibilities toward a tissue-stress interpretation, but it does not choose among them. This framing is intentionally falsifiable: cell-resolved and spatial experiments can test whether high residual tumors contain more NET structures per neutrophil or instead express the program in neighboring compartments.

The persistence of hypoxia and inflammatory coupling also connects this atlas to experimental cancer models. Cathepsin C-driven neutrophil recruitment and NET formation can promote breast-cancer lung metastasis [25], and chronic stress can remodel metastatic niches through neutrophil-dependent mechanisms [26]. Human brain tumors locally activate neutrophils in tissue-specific ways [27], while reciprocal tumor-neutrophil signaling shapes bladder-cancer progression [28]. Most directly relevant to our residual result, hypoxia-driven histone lactylation can reprogram neutrophil function in brain tumors [29]. These studies support a model in which low oxygen, inflammatory signaling and local metabolites change what neutrophils do, not merely how many are present.

Additional work illustrates how the surrounding stroma and systemic environment can feed this state. Stromal amyloid-beta-dependent signaling can induce NETs and modulate tumor growth [30]; serotonin-dependent histone serotonylation and citrullination can drive NETs and liver metastasis [31]; and NETosis has been implicated in radiation and systemic-therapy resistance [32]. These mechanisms make a purely cell-count interpretation inadequate. They also caution against treating the residual as one uniform molecular process: similar transcriptomic coupling may arise through distinct metabolic, cytokine, chromatin or stromal routes in different cancers.

At the cellular level, NET formation encompasses multiple routes of chromatin decondensation, membrane remodeling and granule-protein redistribution [33]. PAD4-dependent chromatin citrullination can participate in a default NET-forming program under some conditions [34], yet neutrophil plasticity in tumors is broader than any single pathway [35]. IL-17-producing gamma-delta T cells can cooperate with neutrophils to promote metastasis [36], and NETosis can be a major driver of thrombosis in immune-mediated vascular disease [37]. The atlas therefore supports an integrated inflammatory-tissue-stress interpretation rather than equating the score with PAD4 activity or with one morphologic NET subtype.

The immunotherapy results are deliberately negative in their strongest test. Productive PD-1 blockade is associated with pre-existing immune-cell positioning and adaptive immune resistance [38], whereas pan-tumor response models integrate variables such as T-cell-inflamed expression and mutational load [39]. T-cell dysfunction/exclusion frameworks [40], interferon-gamma-related expression profiles [41] and tumor mutational burden [42] each capture distinct response dimensions. It is therefore unsurprising that one NET-associated score had AUCs near 0.5 in two small, heterogeneous cohorts. The result is informative because it narrows the translational claim: NET-associated biology may modify immune access or resistance in selected contexts, but is not sufficient by itself to predict response across diseases and drugs.

The phenotype association in IMvigor210 is consistent with a contextual role. Cancer-associated fibroblasts can influence anti-PD-1/PD-L1 response through matrix, cytokine and exclusion programs [43], and physical or functional T-cell exclusion is a central property of immune-privileged tumor niches [44]. Our residual score was negatively coupled to the CAF-endothelial module across cancers, while the raw score varied across inflamed, excluded and desert phenotypes. This apparent tension is useful: raw score, residual score and categorical immune architecture are not interchangeable. A future predictive model should combine NET-associated activation with direct measures of T-cell state, stroma, antigenicity and treatment context.

This context dependence also changes the therapeutic hypothesis. Blocking NET formation would be most rational where active traps are demonstrated and where the surrounding ecology provides a mechanistic route to resistance, such as IL-17 signaling, hypoxia, T-cell exclusion or a stromal barrier. The present atlas does not nominate a single drug or justify indiscriminate DNase, PAD4, CXCR1/2 or complement inhibition. Instead, it supplies a stratification logic for combination studies: enrich for tumors with a high NET-associated score, verify NET structures directly, and then pair the intervention with the dominant contextual liability. A hypoxic NET-associated tumor may require vascular or metabolic normalization; an excluded tumor may require stromal remodeling; an inflamed but resistant tumor may require interruption of a myeloid checkpoint circuit.

The immunothrombosis findings provide a second translational axis. Immunothrombosis was defined as the use of coagulation and platelet pathways as intravascular effectors of innate immunity [45]. Inflammation and thrombosis reinforce each other through endothelial, leukocyte and coagulation circuits [46], and antibodies plus complement can be major thrombotic drivers [47]. The positive raw and quadrant-level immunothrombosis signals are compatible with this architecture. They do not establish clinical thrombosis, because TCGA lacks prospective venous-thromboembolism phenotyping, but they identify tumor types and samples in which molecular and clinical thrombosis measurements would be especially valuable.

That distinction is clinically important. Venous thrombosis varies substantially by cancer type, stage and treatment exposure [48], and validated clinical risk tools such as the Khorana model rely on readily measured patient and tumor variables rather than tumor transcriptomics [49]. The present score should not replace those tools. Its potential role is mechanistic enrichment: identifying an inflammatory, hypoxic and immunothrombotic tissue state for studies that jointly collect pathology, plasma NET markers, complement/coagulation assays and longitudinal thrombotic outcomes. This positioning also fits the expanded hallmarks view of cancer as a disease shaped by interacting host and tumor capabilities [50].

A credible biomarker program should therefore pair four measurement domains rather than optimize the RNA score in isolation. Tumor tissue should establish NET morphology and cellular location; blood should quantify MPO-DNA or citrullinated-histone complexes together with conventional coagulation markers; clinical follow-up should capture thrombosis, metastasis and treatment exposure; and transcriptomic or spatial data should define the hypoxic, stromal and immune context. Discordance among these domains would be scientifically informative. For example, a high transcriptomic score without circulating NET markers may represent a locally confined ecology, whereas high plasma markers without a tumor-tissue signal may reflect systemic inflammation. This paired design is needed to convert the atlas from a mechanistic map into a clinically interpretable assay.

The composition of the published score further argues for this multimodal validation. Its coefficients span chemotaxis, neutrophil activation, tissue factor, adhesion and genes with broader expression across immune or vascular compartments. Several also intersect conceptually or directly with the inflammatory modules tested here. Such overlap is not necessarily a defect: coordinated expression is part of the proposed ecology. It does mean that transcript-to-transcript correlations cannot independently validate NET biology. The strongest future evidence will come from convergence between the score and orthogonal readouts that do not reuse the same genes, including imaging-defined extracellular chromatin, protein-DNA complexes, spatial cell proximity and functional perturbation.

Several limitations bound the inference. First, bulk RNA cannot visualize extracellular chromatin, assign gene expression to specific cells or distinguish vital from lytic NET formation. Second, the neutrophil proxy is curated rather than measured by histology or single-cell deconvolution; residualization removes only its linear component and cannot prove causal independence. Third, the NETs score and several tested modules share inflammatory genes, which can inflate association through biological or mathematical overlap. We retained transparent gene lists and emphasized effect consistency, but orthogonal protein and spatial assays are needed.

Fourth, ESTIMATE and genomic sensitivity inputs covered only BLCA or one available cancer type, respectively; they cannot support pan-cancer robustness claims. Fifth, stage and other covariates were incompletely available, so adjusted Cox models differed in their actual covariate sets and remain exploratory. Sixth, IMvigor210 and GSE91061 differ in cancer type, drug, platform, gene matching and response distribution, and both provide limited power for interaction analyses. Finally, the study lacks experimental perturbation, spatial validation, circulating NET measurements and prospective thrombosis outcomes. These are not cosmetic gaps; they define the validation program required before clinical use.

A practical next step is to prospectively assay the NETs-high/neutrophil-low state. Multiplex immunofluorescence or imaging mass cytometry could quantify citrullinated histones, MPO-DNA complexes, neutrophil localization, T-cell distance and fibrin/platelet deposition in the same tissue. Spatial transcriptomics could test whether the residual program localizes to hypoxic tumor-stromal interfaces. Plasma assays and longitudinal imaging could connect that state to thrombosis and metastasis. Such validation would determine whether the operational ecology identified here marks active NET formation, a permissive microenvironment, or both.

## Conclusions

Reanalysis of a published 19-gene NETs score across 32 cancer types identifies a reproducible, abundance-aware tumor-ecology pattern. The score is strongly linked to neutrophil content, yet hypoxic and TNF-alpha/NF-kB coupling persists after within-cancer neutrophil adjustment and in NETs-high/neutrophil-low tumors. Survival and immunotherapy associations are context dependent. The atlas therefore supports NET-associated transcription as a testable organizer of inflammatory and immunothrombotic ecology, not as direct NET measurement or a universal standalone clinical biomarker.

## List of abbreviations

AUC: area under the receiver operating characteristic curve
BLCA: bladder urothelial carcinoma
CAF: cancer-associated fibroblast
CI: confidence interval
CNV: copy-number variation
FDR: false-discovery rate
GDC: Genomic Data Commons
HR: hazard ratio
ICB: immune checkpoint blockade
MSigDB: Molecular Signatures Database
NETs: neutrophil extracellular traps
NF-kB: nuclear factor kappa B
OS: overall survival
ROC: receiver operating characteristic
TCGA: The Cancer Genome Atlas
TMB: tumor mutational burden

## Declarations

## Ethics approval and consent to participate

Not applicable. This study used publicly available, de-identified datasets and involved no new human-participant recruitment or intervention.

## Consent for publication

Not applicable.

## Availability of data and materials

TCGA data are available through the NCI Genomic Data Commons (https://portal.gdc.cancer.gov/) and UCSC Xena (https://xenabrowser.net/datapages/). GSE91061 is available from NCBI GEO (https://www.ncbi.nlm.nih.gov/geo/query/acc.cgi?acc=GSE91061). IMvigor210 data were analyzed from the processed files described in the original publication. Analysis code, derived summary tables and figure source data are provided as Additional files with this submission.

## Competing interests

The author declares no competing interests.

## Funding

Not applicable.

## Author contributions

Z.L. conceived and designed the study, curated data, developed and verified the analysis workflow, interpreted results, prepared figures, drafted and critically revised the manuscript, and approved the submitted version.

## Acknowledgements

Not applicable.

## Notes

### Competing Interest Statement

The authors have declared no competing interest.

