## Supplementary_material for "Pan-cancer NET-associated immunothrombosis coupling identifies a neutrophil-linked but neutrophil-irreducible tumor ecology": Supplementary_material_Molecular_Cancer_final.pdf

Zhang Lin

#### Supplementary Methods

The ESTIMATE sensitivity analysis used the only locally available matched subset: 396 BLCA tumors. Within BLCA, the standardized NETs score was residualized against the neutrophil proxy plus either standardized stromal score or standardized tumor purity. Spearman correlations with selected pathway modules were recalculated. These results are restricted to BLCA and were not summarized as pan-cancer estimates.

The exploratory genomic analysis harmonized available TMB, mutation-signature proxy and thresholded copy-number burden features to TCGA sample barcodes. Only one cancer type had matched input for each feature. Results are shown to document available evidence and data limitations; they were excluded from the principal novelty claim.

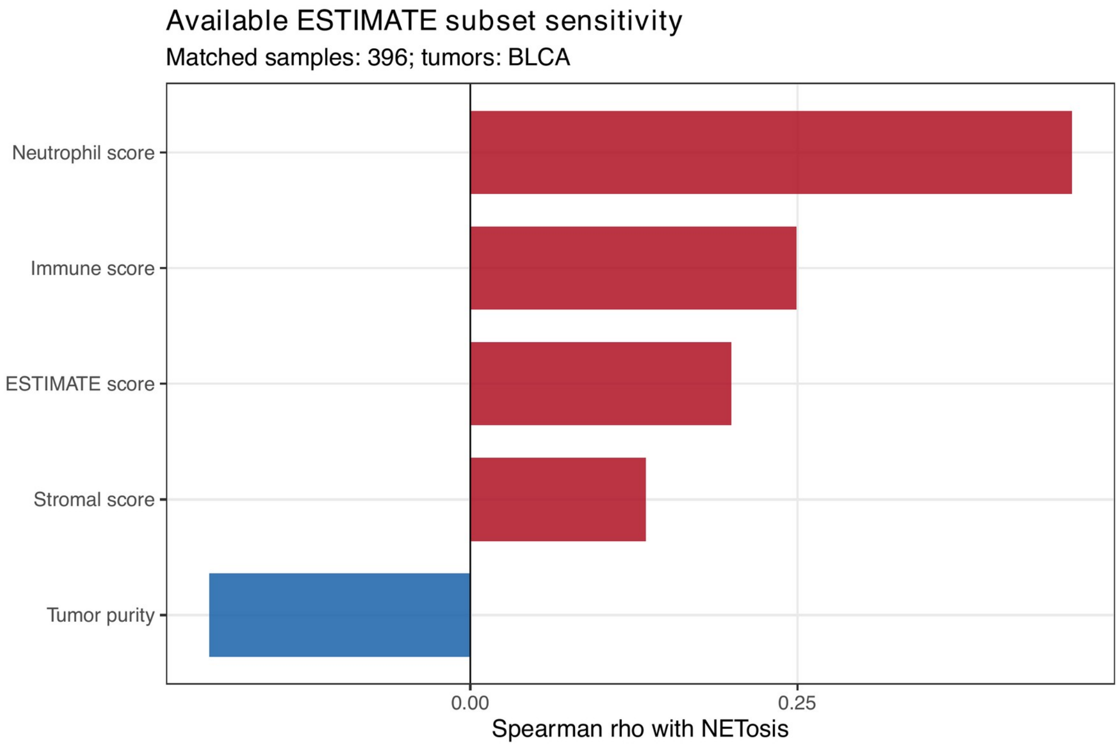


**Figure S1. ESTIMATE sensitivity in the available BLCA subset.** Spearman correlations of the raw NETs score with ESTIMATE, immune, stromal, tumor-purity and neutrophil scores in 396 BLCA tumors. Positive values indicate concordant association; tumor purity is inversely scaled relative to ESTIMATE score. This panel is a BLCA-only sensitivity analysis.

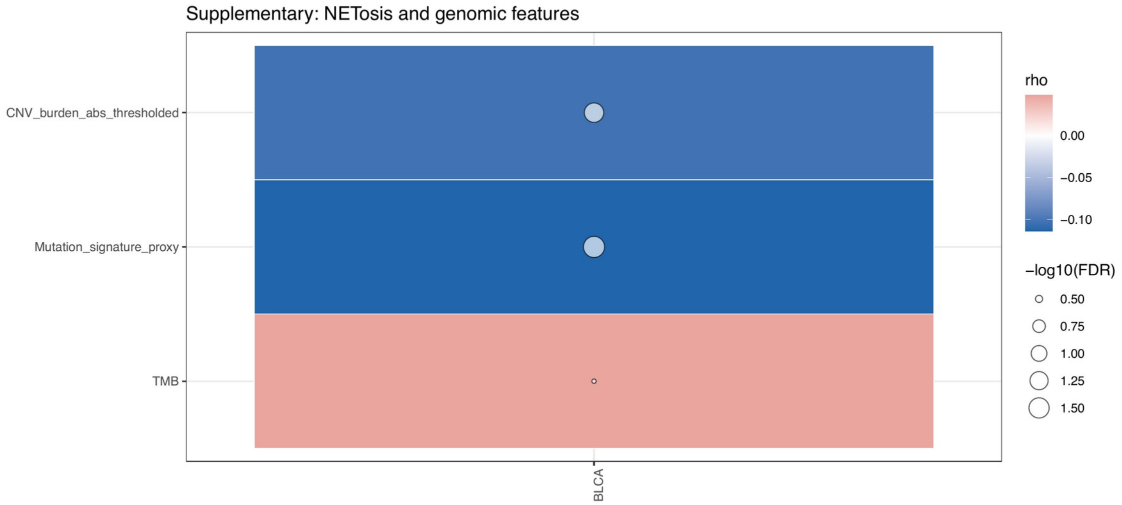


**Figure S2. Exploratory genomic associations in available inputs.** Spearman correlations between the NETs score and TMB, mutation-signature proxy or absolute thresholded CNV burden. Each feature was available in one cancer type only; these estimates do not establish pan-cancer genomic specificity.

### Supplementary Tables

**Table S1. NETs signature gene matching.**

| Dataset | Total genes | Matched | Missing |
|---|---|---|---|
| TCGA | 19 | 19 | None |
| IMvigor210 | 19 | 19 | None |
| GSE91061 | 19 | 17 | CXCL8; VNN3 |

**Table S2. Restricted sensitivity coverage.**

| Analysis | Matched samples | Cancer types | Interpretation |
|---|---|---|---|
| ESTIMATE | 396 | 1 (BLCA) | BLCA-only purity/stromal sensitivity |
| TMB | Available subset | 1 | Exploratory only |
| Mutation signature | Available subset | 1 | Exploratory only |
| Thresholded CNV | Available subset | 1 | Exploratory only |

*Full coefficient, gene-set matching, model-audit, pathway-correlation, quadrant-contrast, immunotherapy and sensitivity tables are supplied as machine-readable TSV files in the supplementary folder and source-data archive.*
